# A low-cost automized anaerobic chamber for long-term growth experiments and sample handling

**DOI:** 10.1101/2020.12.17.423238

**Authors:** Achim J. Herrmann, Michelle M. Gehringer

## Abstract

1

The handling of oxygen sensitive samples and growth of obligate anaerobic organisms requires the stringent exclusion of oxygen, which is omnipresent in our normal atmospheric environment. Anaerobic workstations (aka. Glove boxes) enable the handling of oxygen sensitive samples during complex procedures, or the long-term incubation of anaerobic organisms. Depending on the application requirements, commercial workstations can cost up to 60.000 €. Here we present the complete build instructions for a highly adaptive, Arduino based, anaerobic workstation for microbial cultivation and sample handling, with features normally found only in high cost commercial solutions. This build can automatically regulate humidity, H_2_ levels (as oxygen reductant), log the environmental data and purge the airlock. It is built as compact as possible to allow it to fit into regular growth chambers for full environmental control. In our experiments, oxygen levels during the continuous growth of oxygen producing cyanobacteria, stayed under 0.03 % for 21 days without needing user intervention. The modular Arduino controller allows for the easy incorporation of additional regulation parameters, such as CO_2_ concentration or air pressure. This paper provides researchers with a low cost, entry level workstation for anaerobic sample handling with the flexibility to match their specific experimental needs.

**Specifications table:** *[please fill in right-hand column of the table below]*

|  |  |
| --- | --- |
| Hardware name | *Anaerobic chamber* |
| Subject area | • Biological Sciences (e.g. Microbiology and Biochemistry)<br>• General |
| Hardware type | • Biological sample handling and preparation |
| Open Source License | MIT License |
| Cost of Hardware | ~2000 € |
| Source File Repository | [10.17605/OSF.IO/J84WD](https://doi.org/10.17605/OSF.IO/J84WD) |

## 2 Hardware in context

Handling oxygen sensitive samples can provide a challenge in modern earth’s atmosphere, as oxygen is the second most abundant gas and even trace amounts can lead to the rapid and irreversible oxidation of a sample. Handling oxygen sensitive samples can be further complicated if the sample is alive, like obligate anaerobic organisms, where oxygen and its radical forms acts as a potent toxin, oxidizing important proteins and catalytic metal ligands (Hentges 1996). Cheap solutions like sealed glass tubes or flasks with rubber lids used in combination with syringes, produce a lot of waste and their use is limited if complex sample handling is required. This hardware offers flexibility in manual sample processing, in an anaerobic environment, in an affordable, programmable anaerobic workstation.

## 3 Hardware description

This paper describes the construction of a low-cost alternative to commercial anaerobic chamber with advanced features usually only found in high cost, automated anaerobic chambers. Barebone anaerobic workstations with glove ports can easily cost more than 10.000 €, even without “comfort” functions like an automated airlock or automatic maintenance of an anaerobic atmosphere. Advanced features like humidity or CO2 control quickly add 5,000 to 10,000 € to the cost, depending on the feature and company in question. Here we describe the complete build instructions to a cheap, Arduino base, automated anaerobic workstation, which can be used in the cultivation of bacteria or for general sample handling. In contrast to other DIY solutions, the automated control allows for long term experiments or storage of samples without user input. Additional features like CO2 regulation or pressure control can be added as needed per experimental requirement, even after the final installation.

Advantages of this design:

- automatic regulation of humidity and H_2_ (as oxygen reductant) concentration
- Logger for monitoring environmental conditions during the experiments
- Automatic air-lock purging
- Easily expandable for individual requirements like controlling the CO_2_ concentration
- Compact design allows for the integration into growth chambers/cell incubators and/or minimal impact on available lab space

Possible use cases:

- Growth of anaerobic organisms
- Growth of oxygen producers like cyanobacteria under anaerobic conditions
- Handling and storage of oxygen sensitive samples
- Radioactive labeling experiments under anaerobic conditions

## 4 Design files

### Design Files Summary

| Design file name | File type | Open source license | Location of the file |
| --- | --- | --- | --- |
| Mainbody | CAD .fcstd | MIT | [10.17605/OSF.IO/J84WD](https://doi.org/10.17605/OSF.IO/J84WD) |
| FrontDoor | CAD .fcstd | MIT |  |
| GloveGasket | CAD .fcstd | MIT |  |
| GloveLid | CAD .fcstd | MIT |  |
| AirlockTop | CAD .fcstd | MIT |  |
| GasPortAirlock | CAD .fcstd | MIT |  |
| GasPortTop | CAD .fcstd | MIT |  |
| FanHolder | CAD .fcstd | MIT |  |
| HolderSensor | CAD .fcstd | MIT |  |
| LSensorHolder | CAD .fcstd | MIT |  |
| ScrewBacking25 | CAD .fcstd | MIT |  |
| ScrewBackingFlat | CAD .fcstd | MIT |  |
| ChamberPipeInlet | CAD .fcstd | MIT |  |
| ChamberPipeOutlet | CAD .fcstd | MIT |  |
| OverpressurePipe | CAD .fcstd | MIT |  |
| CompleteAssembly | CAD .fcstd | MIT |  |
| ArduinoCad | Electronic .dsn | MIT |  |
| SS3_eng | Software .ino | MIT |  |
| SS3_eng_setup | Software .ino | MIT |  |

“Mainbody” describes the body onto which all parts are assembled. It was assembled from multiple slabs of PMMA, which were cut and drilled, before being glued together. In order to reduce costs, the bottom, sides and back plate could be made from a cheaper material, as only the front and top part needs to be translucent in order to see inside.

“FrontDoor” describes the front door of the airlock, which seals the box hermetically. The groove on the inside should be fitted with the round cord to ensure an airtight seal.

“GloveGasket” describes the gasket to secure the gloves in place during operation and ensure an airtight seal. The groove on the inside should be fitted with the round cord to ensure an airtight seal. For an easier installation of the gloves inside the gaskets, clear sticky tape can be used to hold the gloves in place.

“GloveLid” describes a cover for the glove holes, if the workstation is not in use, to prevent the gloves from sticking out due to the internal over pressure.

“AirlockTop” describes the top cover of the airlock. Holes for the toggle fastener are not described, as they are part dependent, and have to be adjusted for a tight fit with the seal strip.

“GasPortAirlock” describes a block with screw fittings for the airlock gas in- and outlet, as well as the main chamber gas outlet. It needs to be glued in the Gas outlet bay (<u>Fig. 4</u>).

**Fig 1:**
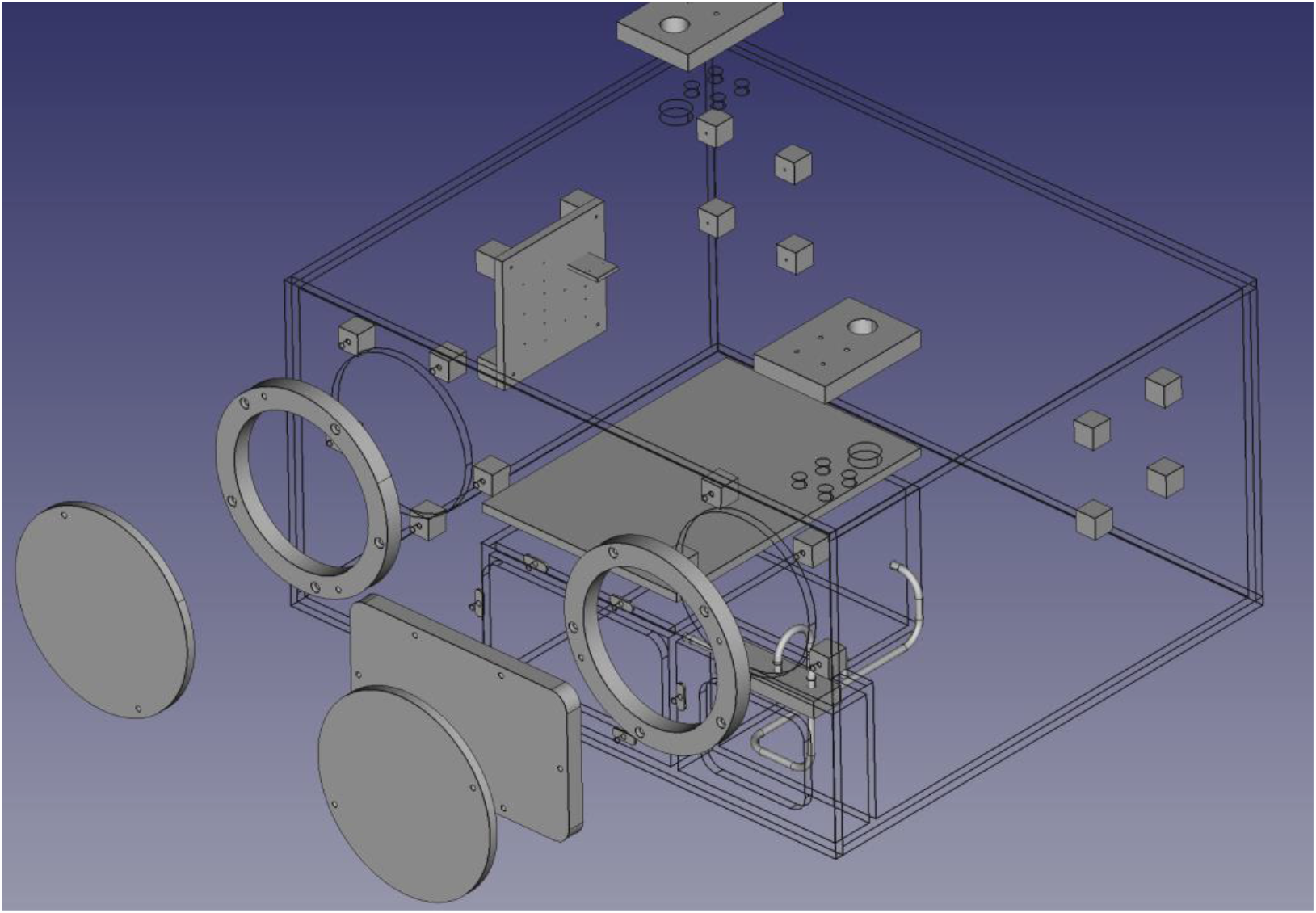
Overview of all the parts assembled from the file “CompleteAssembly”.

**Fig 2:**
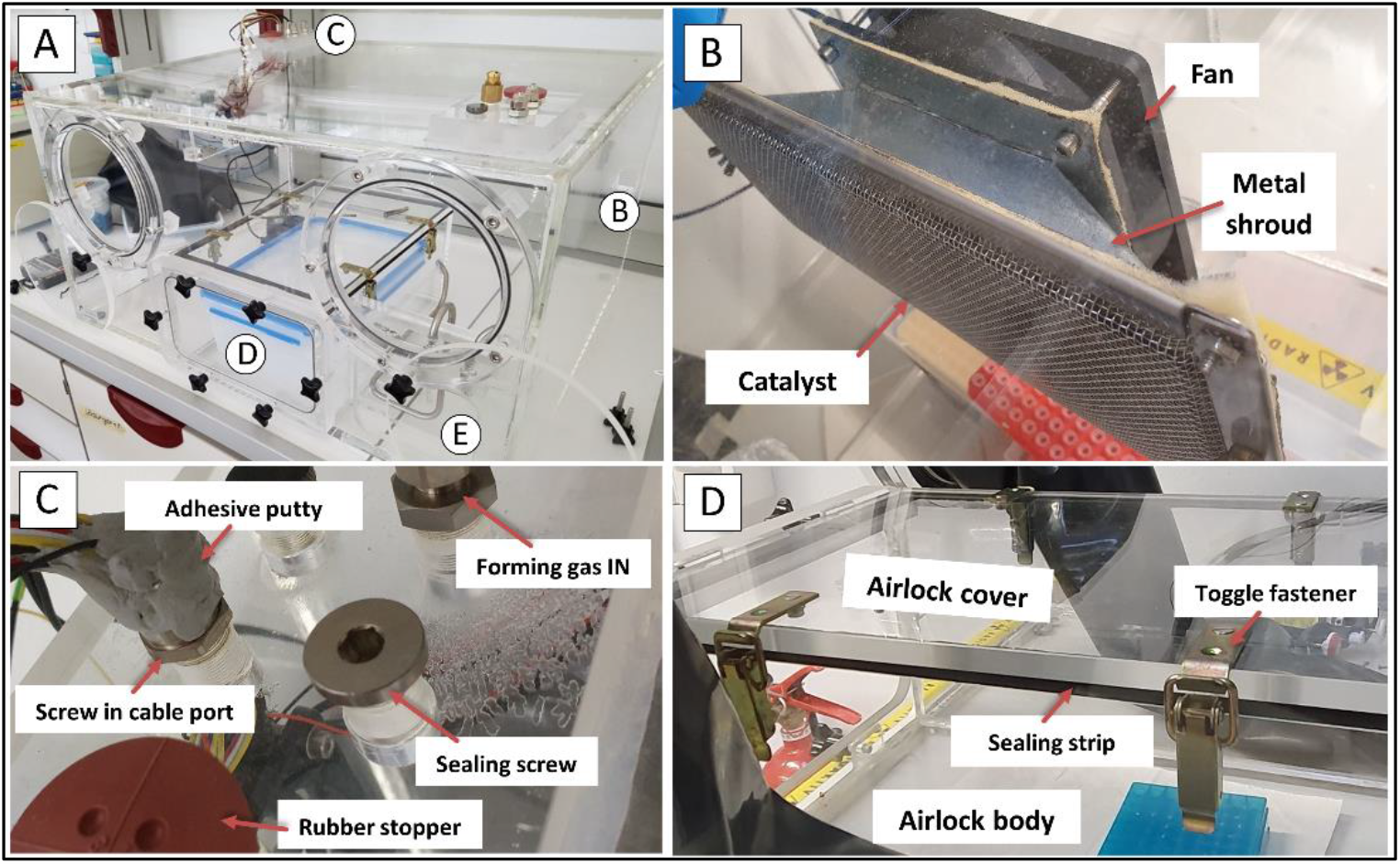
A: Completely assembled PMMA body of the anaerobic chamber with round nominators for in detail pictures. B: Final catalyst assembly with a metal shroud connecting it to one of the fans screwed to the fan holders. C: Cable entry port in detail. D: Airlock assembly with cover, sealing strip and closed toggle fastener.

**Fig 3:**
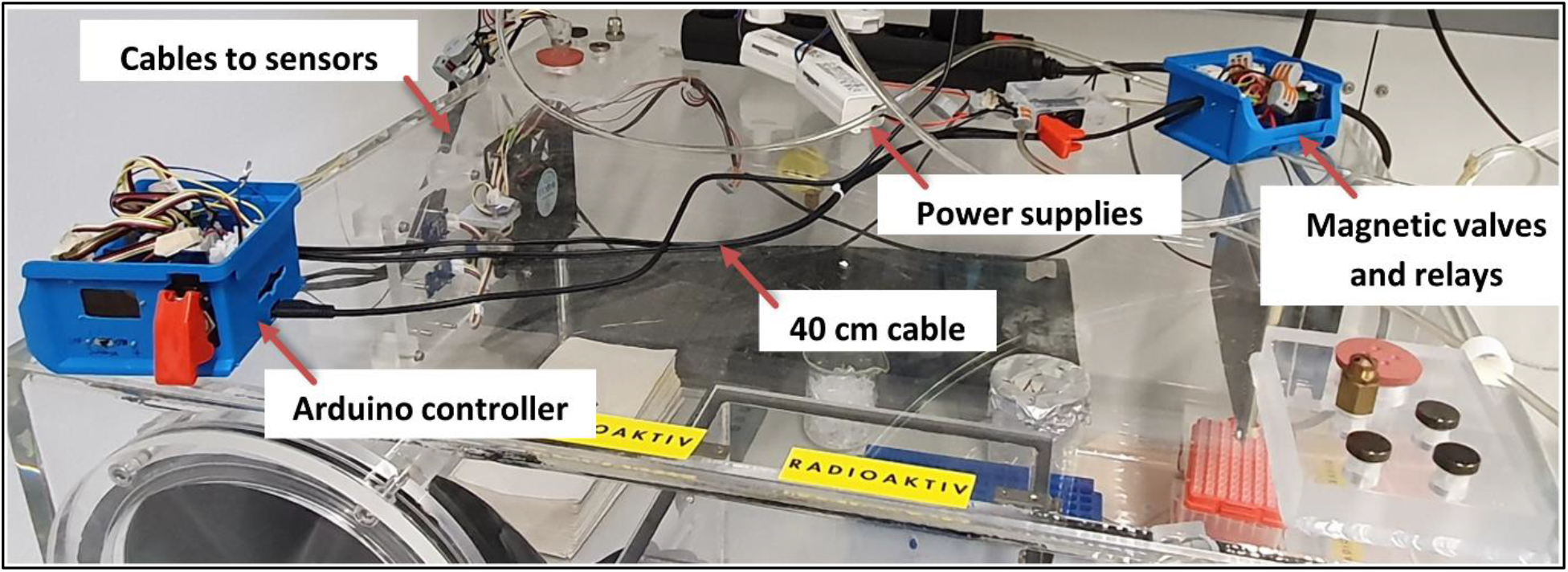
Spatial separation between the Arduino controller and the magnetic gas valves and power supply units to prevent interference.

**Fig 4:**
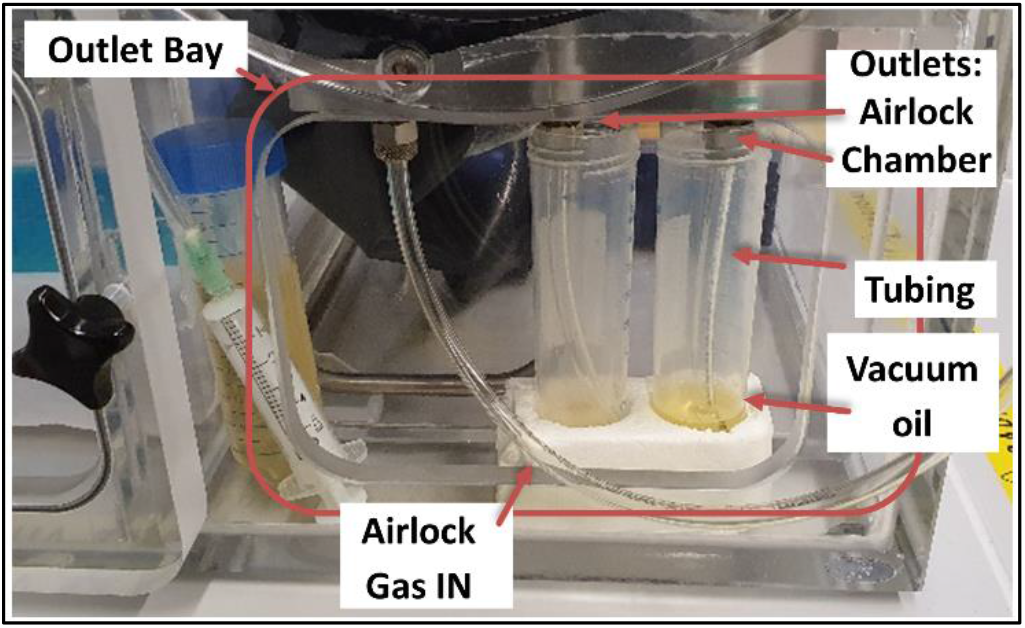
Gas outlet bay with 50 ml test tubes filled with vacuum oil filed just above the tube opening.

“GasPortTop” describes a block with screw fittings for the gas inlets and cable port. The biggest hole is for a Rubber stopper diaphragm, if the drawing of gas with a syringe is needed.

“FanHolder” describes single blocks for attachment of the pc fans and the sensor holder. These need to be glued in sets of four, with the required spacing inside the main body, to ensure a constant removal of oxygen.

“HolderSensor” describes the mounting plate for up to five Grove sensor shields and the “LSensorHolder” for an upright mounting of the light sensor. Needs to be mounted on one set of “FanHolder”.

“ScrewBacking25/-Flat” describe pieces of PMMA which cover the screw holes in the main body for an airtight compartment.

“ChamberPipeInlet/-Outlet” describe the pipes needed to connect the airlock in and outlet to the “GasPortAirlock”.

“OverpressurePipe” describes the pipe connected to the “GasPortAirlock”, where pressure from the main box is vented. It is needed to prevent the gloves from accidentally obstructing the gas outlet.

“CompleteAssembly” shows the complete assembly of all the parts so far described (<u>Fig. 1</u>).

“ArduinoCad” shows the layout of the electric connections of the sensors to the Arduino controller, as well as the power supply and relay cabling. It was created using TinyCad version 3.00.02.

“SS3_eng” is the main software running on the Arduino controller during operation of the anaerobic workstation.

“SS3_eng_setup” is used during the initial sensor and timing setup and for verification of sensors functionality (see <u>6</u>).

## 5 Bill of Materials

The bill of materials is uploaded to the open science framework at <u>10.17605/OSF.IO/J84WD</u>.

## 6 Build Instructions

Assemble the PMMA body of the anaerobic box (Fig. 2A) as per construction file (CompleteAssembly). Be sure to install all internal parts before sealing the cover plate, especially the airlock top plate (AirlockTop), as it is too big to be inserted through the airlock afterwards. The airlock cover plate must be connected to the airlock main body with four toggle fasteners (Fig. 2D). The boreholes for the toggle fasteners are not specified in the blueprints and should be adjusted to ensure a tight fit with the underlying sealing strip if the toggle fasteners are engaged. The sealing strip is attached to the top of the airlock walls.

Mount the fans on the fan holders and attach the catalyst (StakPak) in front of one of the fans (Fig. 2B). In our case, this was achieved by a folded metal shroud made of 2 mm steel. As a cost saving measure, a catalyst can be easily made from scratch: 10 to 20 g of Palladium on Carbon (5%) pellets (2-3mm) should be encased between two layers of fine stainless-steel wire mesh. The sides can then be sealed by folding them and stapling them shut. Do not use flammable substances in the construction as the catalyst can become hot during operation and could melt plastic or char cardboard.

Fan power and connection cables are passed through a screw-in cable port and the port sealed airtight with either adhesive putty or epoxy resin (Fig. 2C). Connect all sensor cables to the Arduino as described in the “Arduino CAD” file.

The Arduino controller must be electronically shielded or spatially separated from the magnetic gas valves and their power source, as their operation can create strong magnetic fields and hence system instability. In our case, this was achieved by placing the Arduino in one case made from a plastic storage box, while the gas valves with their relays were placed in a second plastic box and connected with 40 cm of wiring (Fig. 3).

The gas sensors for H_2_ need to be calibrated using the Arduino Program “SS3_eng_setup”. Open the serial monitor and note the R_0_ value of the H_2_ sensor in ambient air. This value needs to be adjusted in the main program “SS3_eng”. Additionally, the other sensors should be checked for their functionality in the setup. The setup file also sets the Real Time Chip. Adjust the files in void setup in SS3_eng_setup” according to your current date and time:

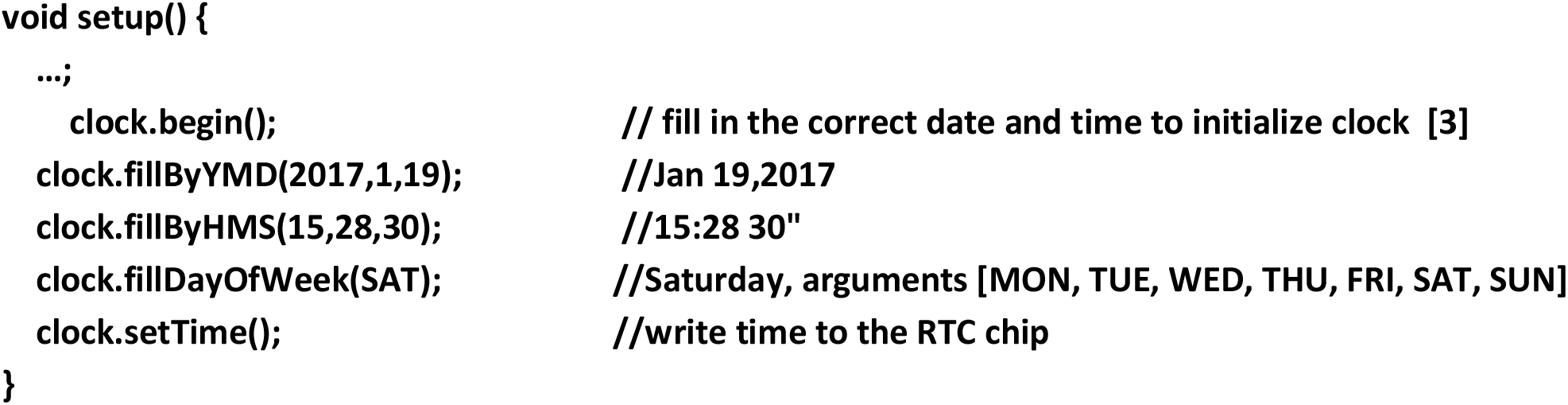

The above process sets the time to the specified values every time the code is run and should be deleted after the initial setup or the displayed time may be inaccurate.

After measuring the R_0_ value, adjust the lines in the main program “SS3_eng”:

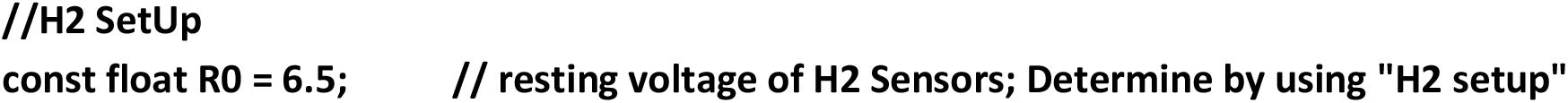

Further information on the function, principal and application of the utilized sensors can be found on: https://wiki.seeedstudio.com

### Safety instructions

The anaerobic workstation uses non-toxic, asphyxiating gases and should not be operated in confined spaces without ventilation. If forming gas with ≤ 5% H_2_ is used, there is no explosive risk, as the minimal flammability limit cannot be reached by mixing forming gas with ambient air. The risk of electric shock is minimal as the maximal voltage used is only 24V.

## 7 Operation Instructions

The ends of the chamber/airlock gas outlet tubes must be submerged in water or oil to prevent the inflow of ambient air. This can be achieved either by placing the end of a short section of stiff tubing inside a liquid filled reaction tube or similarly sized vessel inside the outlet bay or running the tubing to an external gas washing flask or similar device. Care should be taken to not completely fill the vessels, as the immersion depth of the tubing correlates to the maximum internal overpressure during gassing operations. High internal pressure can make the gloves rather stiff and unpleasant to handle while also posing the risk of an outward spillage of oil/water during a rapid insertion of the gloves. Likewise, a rapid retraction of the gloves poses the risk of sucking in liquid and ambient air contaminating the internal atmosphere/ surroundings. Therefore, care should be taken while inserting and removing one’s arms and, if necessary, gas can be manually inserted by activating the manual gas injection switch to alleviate an eventual negative pressure. Tip: If the liquid level in the container, which houses the outlet from the airlock, is lower than in the chamber outlet, less ambient air will enter the main chamber during an airlock cycle.

N_2_ can be used in certain steps as a cost saving measure. All steps describing the usage of N_2_ gas can also be performed with forming gas if no N_2_ is available.

### 7.1 Initial Setup

1. Make sure all tubes are connected, all unused screw ports are covered by sealing screws and all internal sensors are working. Fill up the outlet vessels with oil/water to slightly cover the tube endings (∼5 mm).
2. All necessary equipment like pipettes, test tube holders e.g. should be inserted before making the chamber anaerobic, as this can save time otherwise spent on unnecessary airlock cycles.
3. (Optional)Flush the box with N_2_ gas for 20 min to bring down the O_2_ levels and save forming gas.
4. Switch on the controller with the connected forming gas. Forming gas injection will start to remove the remaining O_2_. Humidity should rise as O_2_ reacts with H_2_ to form H_2_O. This exothermic reaction can heat up the catalyst and damage delicate equipment if they are in direct contact.
5. The controller should auto adjust the injections of forming gas and N_2_ until an atmosphere is established within the parameters set for humidity and enough H_2_ gas to remove all O_2_ present in the anaerobic chamber. As the frequency of gas injections during setup is most likely over the safe operating threshold during normal operation, the gas override switch should be switched to ON <u>(7.2</u>).

### 7.2 User interface

The OLED display (Fig. 5) shows all relevant environmental conditions during the operation:

**Fig 5:**
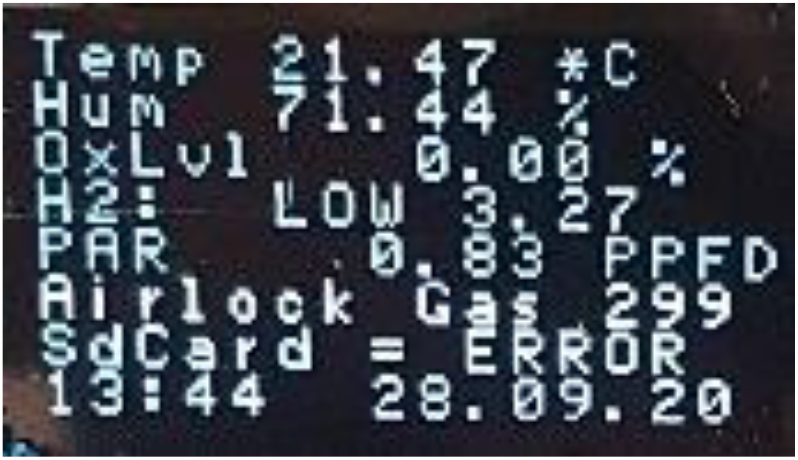
User interface on OLED display.

1. Temperature in °C
2. Relative Humidity in % saturation at the temperature in line 1
3. Oxygen levels in % with an error of 0.03%
4. Indication of the H_2_ level in the chamber with:
  a. NONE ∼ H2 < 1%
  b. LOW ∼ H2 < 3%
  c. NORM ∼ H2 >3% The number is displays the Rs (Reading Sensor)/ R_0_ (Reading Zero) ratio, NOT H_2_ in % (Fig. 6), and is a rough indication of the presence of hydrogen in the atmosphere. As the sensor can also detect alcohols, alcohol-based disinfectants should not be used inside of the glove box, otherwise the auto injection of forming gas will not work.

**Fig 6:**
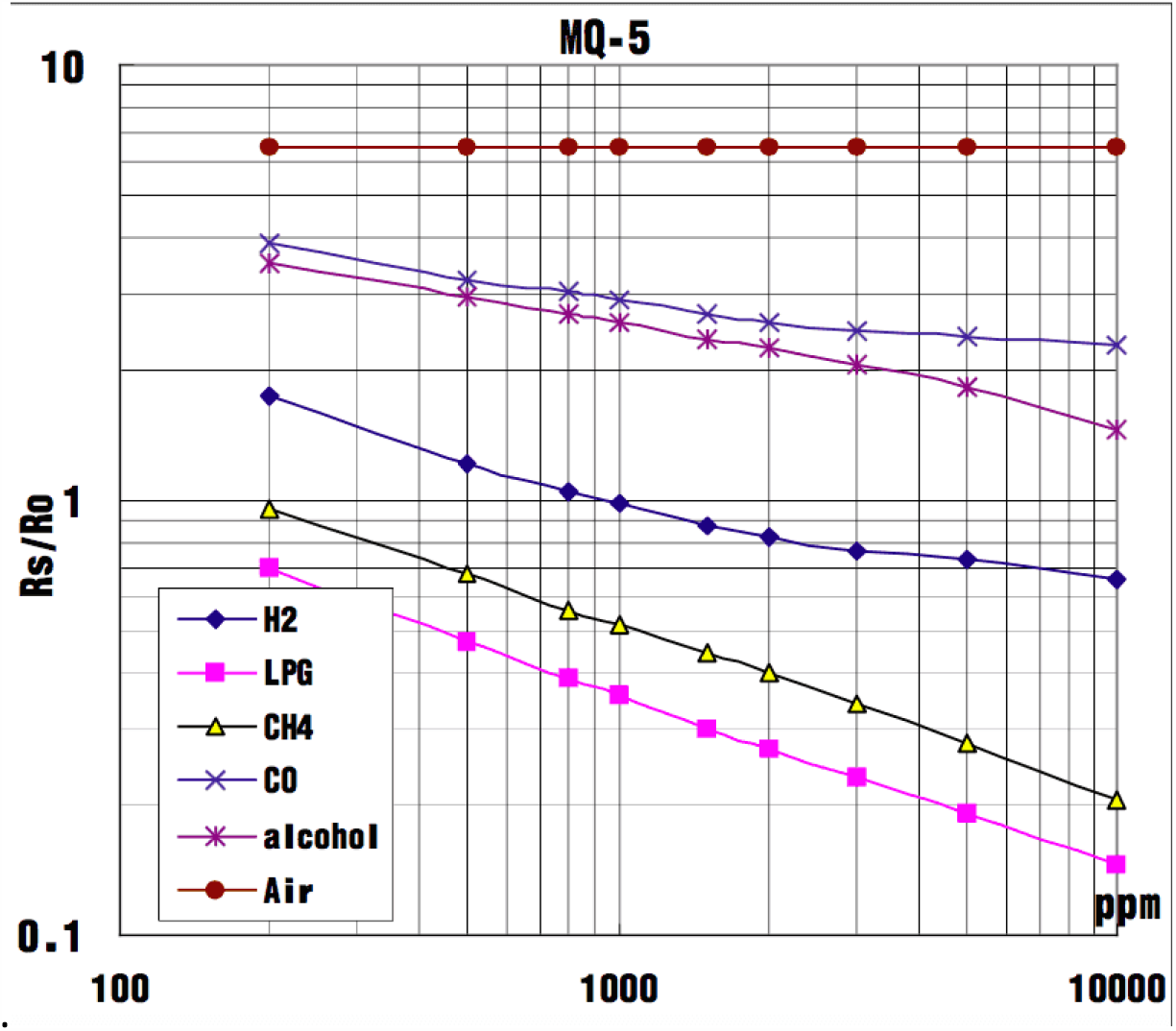
R_S_ (Reading Sensor)/ R_0_ (Reading Zero) value gives an indication of the gas levels in the atmosphere. Note that the sensor will react to other gases in the atmosphere, which should be taken into account during the operation of the automated gassing system (https://wiki.seeedstudio.com/Grove-Gas_Sensor-MQ5/; 13.10.2020)

**Fig 7:**
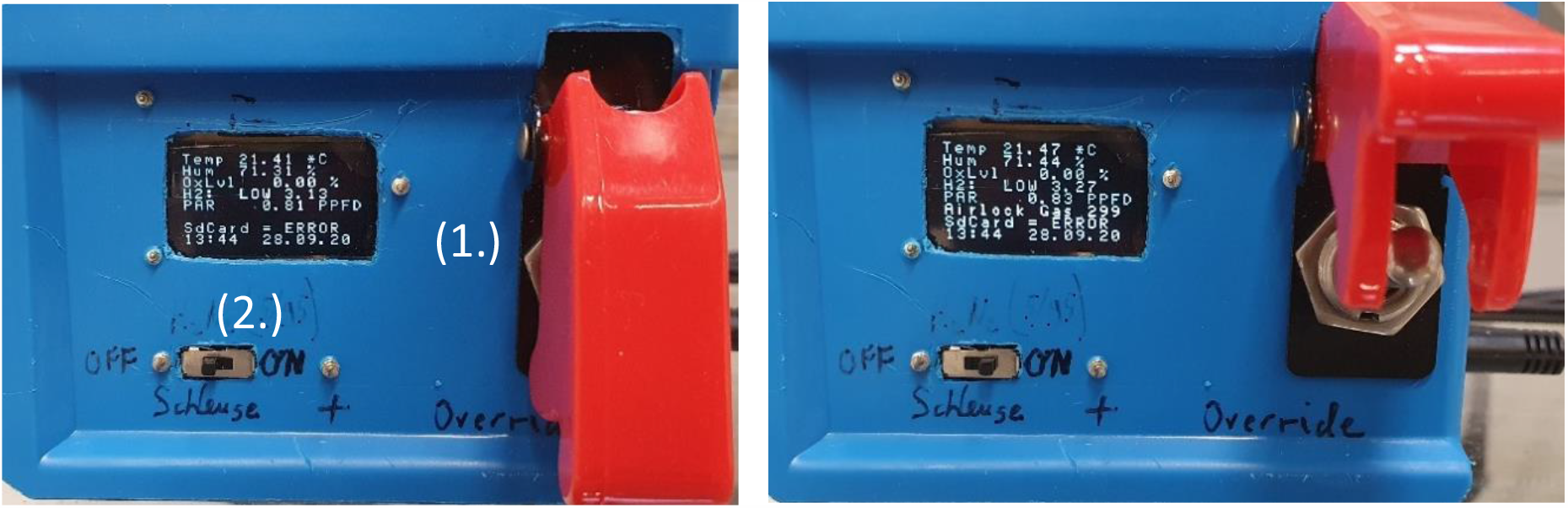
Operator control interface with the Killswitch (1) and Slide Switch (2) disengaged (left) and engaged (right). Note the change in the OLED display line 6.

**Fig 8:**
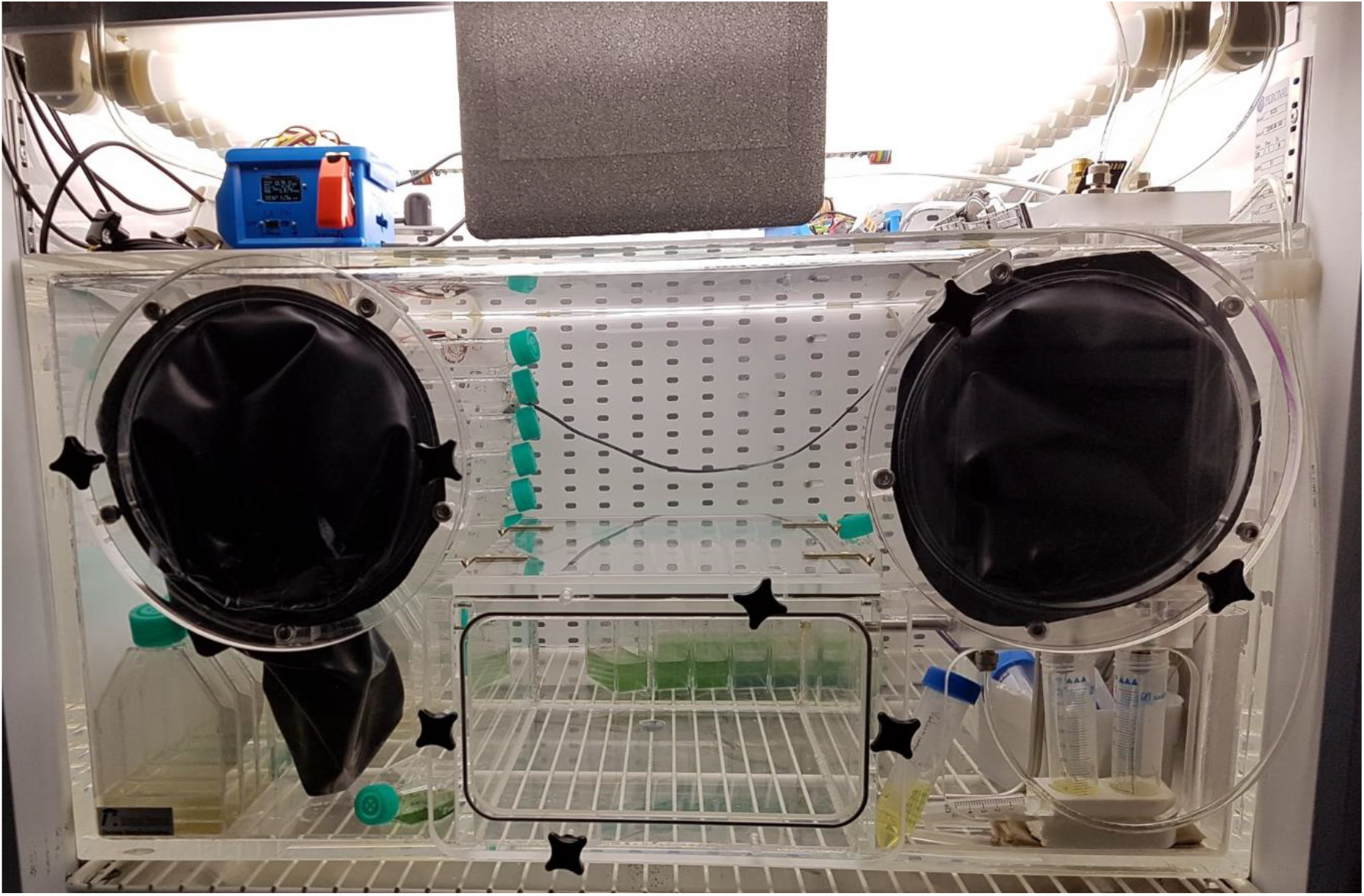
Anaerobic chamber fully functional and wired inside a culture chamber with growing cyanobacterial cultures in ventilated culture flasks.

**Fig 9:**
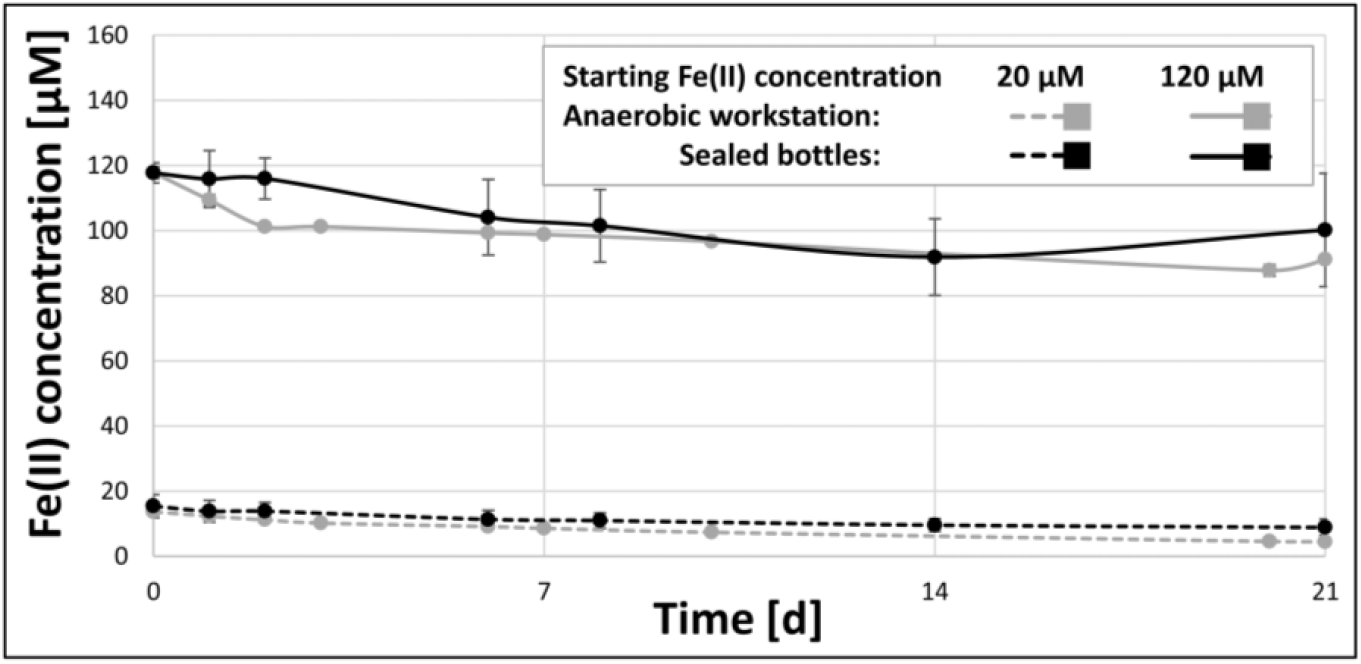
The Fe(II) concentrations of growth media over the course of 21 days, if either incubated inside the here described anaerobic chamber (grey), or in sealed anaerobic bottles (black). The starting Fe(II) concentrations were either set at 20 µM (dotted lines) or 120 µM (solid lines).
5. PAR in PPFD. Only works if the light sensor is calibrated, otherwise just an indicator of the current day-night / light-dark cycle
6. Status of the automatic gas injection
  a. GASSING ERROR: The auto injection was disabled because gas was injected too often in a short interval, hinting at a leak or sensor malfunction. To reset this message and enable auto injection flip the OVERRIDE switch to ON and return to OFF.
  b. GAS OVERRIDE: Displays the status of the OVERRIDE switch as ON.
  c. AIRLOCK GAS = n: Automatic gassing of the airlock with n seconds remaining
7. Status of the SDCard of the automated saving of data
  a. ERROR = SD Card is not mounted, either no SD Card is inserted or it could not be mounted
  b. OK = SD Card is mounted and data gets saved every minute
8. Time and date

The controller has two operational modes, which can be controlled by switching the Killswitch (1) and Sliding Switch (2) to the corresponding positions.

1. Killswitch (1) is OFF:
  a. Slide Switch (2) is OFF: Emergency Stop Function is active: the controller checks if 5 consecutive gas injections were triggered within 10 seconds of each other (indication of a leakage or an empty gas bottle) and stops further gas injection without user input.
  b. Slide Switch (2) is ON: Manually injects forming gas. Useful to prevent negative pressure while operating the gloves.
2. Killswitch (1) is ON:
  a. Slide Switch (2) is OFF: Display shows GAS OVERRIDE. Resets the emergency Stop Function and disables it while the Killswitch (1) is engaged.
  b. Slide Switch (2) is ON: Floods the airlock chamber with N_2_ for 300 seconds (can be adjusted in program file) and displays the remaining time on the display. Gassing can be stopped by disengaging any off the input switches (1 or 2) to OFF. In order to minimize the inflow of ambient air from the airlock chamber into the main body, a tight fit of the inner airlock top cover must be ensured. During the first minute of gassing, the airlock front door can also be slightly opened for venting in order to prevent any gas from leaking into the chamber.

### 7.3 Cleaning and maintenance

Avoid using solvents during the cleaning and disinfection of the anaerobic chamber as the MQ5 sensor detects not only H_2_ but also hydrogen in hydrocarbons. It may give a false indication of H_2_ levels, if volatile solvents like ethanol or isopropanol are present in the anaerobic chambers atmosphere. Solvents can also lead to cracking or cloudiness of the PMMA material and adsorb onto the activated charcoal catalyst, reducing its activity. In order to remove adsorbed solvents from the catalyst, it can be baked overnight at 200°C (Caution: depending on the quantities adsorbed on the catalyst this can pose an explosion hazard!!!). Therefore, only use water and mild detergents during cleaning operations if possible.

Care should be taken during the incubation of organisms that produce or require SO_2_/H_2_S or CO as these gases can lead to a poisoning of the palladium catalyst (Albers et al. 2001).

The oxygen sensor detects oxygen by slowly reacting with O_2_, producing a current, and lasts for ∼ 2 years under ambient oxygen levels. In order to maximize the sensor lifetime, it should be stored in an anaerobic environment, if the anaerobic chamber is not currently in use.

## 8 Validation and Characterization

The anaerobic chamber described in this paper was successfully used in studying the effects of Fe(II) on cyanobacteria. To do so, the anaerobic workstation was inserted inside a Percival culture chamber (E-22L), which controlled lighting, temperature and CO_2_ levels via an external sensor. In order to test for the effect of photooxidation of Fe(II), or the buildup of oxygen inside the anaerobic chamber, control cultures without cyanobacteria were set up and Fe(II) measured throughout the whole experiment by means of the colorimetric ferrozine assay. The whole experiment was then repeated inside discrete, hermetically sealed, anaerobic bottles (which were also set up inside the anaerobic chamber) to test for the effects of O_2_ buildup on cyanobacterial growth.

The Fe(II) measurements of the controls showed that the anaerobic box maintained an anerobic atmosphere throughout the whole experiment just as well as the individually sealed bottles. Under aerobic conditions, the complete oxidation of Fe(II) would have been observed within 20 minutes. The slight drop in Fe(II) concentration over the whole length of the experiments is most likely the result of photo-oxidation, as the media was constantly exposed to the light needed for the growth of the cyanobacteria. This result clearly demonstrates the efficiency of the anaerobic workstation, as the control cultures in ventilated flasks were incubated at the same time as the cyanobacterial cultures inside the anaerobic workstation. Despite the cultures producing ∼ 170 ml of pure O_2_ per day near the end of the experiment, the Fe(II) in the control flasks was not oxidized. Without the automated injection of H_2_ and the regulation of humidity, this high influx of oxygen would have required manual user intervention at least every second day. An open bottle placed inside the anaerobic chamber during the experiments with media containing resazurin (20 mg × l^-1^), a commonly used indicator for anaerobic conditions, also showed no signs of oxygen in measurable quantities.

The anaerobic box was also used for the study of phosphorylation of redox sensitive proteins using radioactive ^32^P, hence the radioactive stickers in Fig. 3. The thick PMMA body of the anaerobic chamber offers better protection against beta-radiation then the thin plastic used in anaerobic tents, while the compact interior also allows for an easier clean-up after the end of the labeling experiments.

## 9 Acknowledgements

This project was funded by the German Research Foundation, DFG, Grant number: GE2558/3-1 & GE2558/4-1 awarded to MMG. The authors wish to thank the employees of the metal workshop (Technische Universität Kaiserslautern), especially Mr. Christian Rahm, for assistance in the planning and the assembly of the PMMA body.

## 10 Declaration of interest

Declarations of interest: none

## 11 Human and animal rights

No human or animal studies were conducted in this work.

## Notes

https://doi.org/10.17605/OSF.IO/J84WD

### Competing Interest Statement

The authors have declared no competing interest.

